# Antenatal microbial colonisation of mammalian gut

**DOI:** 10.1101/236190

**Authors:** Elisa Borghi, Valentina Massa, Marco Severgnini, Grazia Fazio, Laura Avagliano, Elena Menegola, Gaetano Bulfamante, Giulia Morace, Francesca Borgo

## Abstract

The widely accepted dogma of intrauterine sterility and initial colonisation of the newborn during birth has been blurred by recent observations of microbial presence in meconium, placenta and amniotic fluid. Given the importance of a maternal-derived *in utero* infant seeding, it is crucial to exclude potential environmental or procedural contaminations, and to assess foetal colonisation before parturition. To ascertain antenatal microbial colonisation in mammals, we analysed sterilely collected intestinal tissues from rodent foetuses in parallel with experimental controls, and tissues from autoptic human foetuses. Next generation sequencing (NGS) showed the presence of pioneer microbes in both rat and human intestines, as well as in rodent placentas and amniotic fluids. Live microbes were isolated from culture-dependent analyses from homogenized rat foetal intestines.

Microbial communities showed foetus- and dam-dependent clustering, confirming the high interindividual variability of microbiota even in the antenatal period. Fluorescent *in situ* hybridisation analysis confirmed the microbes’ existence in the lumen of the developing gut.

These findings have vast implications for an emerging field of enhancing the management of healthy pregnancies, and for understanding how the infant microbiome starts and it is thus shaped.

## INTRODUCTION

Foetus, amniotic fluid and chorioamnion tissue have long been considered sterile until birth or rupture of the amniotic sac. However, recent evidence shows that the intrauterine sac environment harbours a diversity of microorganisms even in physiological pregnancies [1-3], contradicting the long-standing dogma of “womb sterility” [4]. Indeed, the cultivable genera *Enterococcus, Streptococcus, Staphylococcus,* and *Propionibacterium* have been isolated from umbilical cord blood of healthy neonates born by caesarean section [5]. Within the abundant literature on human placenta microbiome composition [3,6,7], a recent study combining 16S ribosomal DNA-based and whole-genome shotgun metagenomic analyses, showed the presence of a unique placental microbiota, strongly resembling the maternal oral bacteria, with the dominance of *Firmicutes, Tenericutes, Proteobacteria, Bacteroidetes,* and *Fusobacteria* phyla [3]. In addition, amniotic fluid and placenta were found to have similar microbial communities, consistent across individuals [6]. Lactic acid bacteria and enteric bacteria have been reported in meconium collected after birth [8], and a certain degree of similarity has been demonstrated between meconium and amniotic fluid [6], probably related to liquid swallowing by the foetus during pregnancy.

All these data suggest but not prove that humans might come in to contact with bacteria before birth, and, depending on the time of gestation and the type of bacteria that first seed the foetus, this antenatal colonization might have important physiological and clinical consequences. Indeed, microbes, either true pioneer or transient species, could expose the developing foetus to a diverse array of antigens [9] that educate the foetal immune system toward tolerance and participate in the full development of the gut-associated lymphoid tissue [10].

At the same time, toxins and viable microorganisms, through active and passive transport from maternal circulation to the placenta could gain direct entry into foetal circulation, eliciting infective and/or inflammatory processes [11].

Despite these recent advances in the field, a conclusive analysis of antenatal microbial colonisation has not been reported [12], leaving a gap in the knowledge of this important developmental process. The present study is aimed at ascertaining antenatal microbial colonisation of mammalian foetal intestinal tissues. Due to the vast medical and scientific consequences of a “colonised womb”, it is crucial to exclude potential environmental or procedural contamination. To address this issue, a rodent animal model was used to allow sterile experimental conditions. This was compared to human gut samples from foetal autopsies that were studied using a 16S rRNA amplicon-based NGS approach, validated by *in situ* detection.

## RESULTS

### *Microbial species are identifiable and viable in rodent foetuses* in utero

We collected, under sterile conditions as described in the method section, intestine, placenta and amniotic fluid from five rat foetuses: three foetuses (numbered 1-3) from one dam (dam A) and two (numbered 4-5) from the other (dam B). The tissues were analysed by 16S rRNA sequencing.

An average of 259,465 reads were obtained per sample, giving a total of 4,670,364 reads overall. Paired-end reads generated from the original DNA fragments using Illumina MiSeq Next Generation Sequencing were merged and quality-filtered producing a total of 1,560,296 sequence tags from the gut samples, and 982,017 and 900,070 from placentas and amniotic fluids, respectively.

9 different bacterial phyla were identified in rat foetal samples following negative control subtraction. The most represented phyla (Fig. 1A), using a cut-off applied of a relative abundance greater than 1% in at least one experimental group, were *Firmicutes* (mean relative abundance ± sd, 57.0±8.6), *Bacteroidetes* (23.7±8.7), *Actinobacteria* (10.3±8.4), *Proteobacteria* (5.0±2.1), and *Verrucomicrobia* (2.8±1.9).

**Figure 1.**
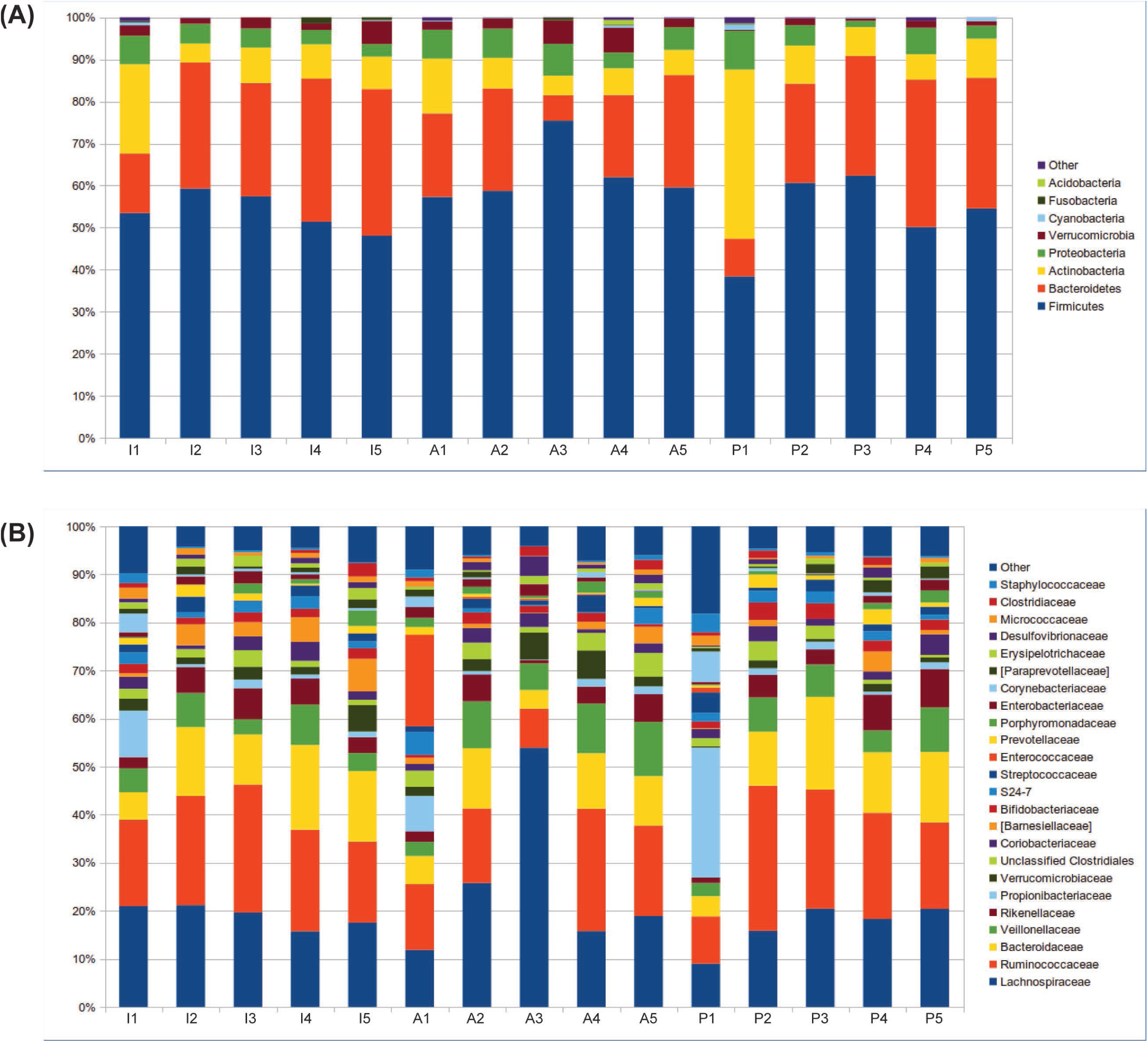
Pioneer microbiota in the developing rodent gut. Bar charts representing the relative abundance of 5 foetal intestines (I1-I5), amniotic fluids (A1-A5) and placentas (P1-P5). The figure shows relative abundance of bacterial **(A)** phyla and **(B)** families.

The most abundant families (Fig. 1B) were: *Ruminococcaceae* (20.9±7.6), *Lachnospiraceae* (20.5±9.3), *Bacteroidaceae* (11.4±4.4), *Veillonellaceae* (5.9±3), *Rikenellaceae* (4.2±2.3), and *Propionibacteriaceae* (3.5±6.3).

To assess whether identified bacteria were viable, two different experiments were performed using a pool of three dissected foetal intestines, sterilely collected and stored at -80°C. Following a growth enrichment step, Gram stain was performed and few dispersed bacteria were detected. In the second experiment, we performed a further step of enrichment in anaerobic blood culture bottle prior subculturing. The bottle turned out positive on day 4, revealing the presence of live microbes. Colony picking and MALDI-TOF processing on the second experiment cultured bacteria allowed for the identification of *Propionibacterium* spp..

### Microbial community is characteristic of foetuses and dams

In order to understand the main determinants constituting the microbial diversity, we evaluated the differences amongst the samples on both richness and composition. Tissues (i.e. intestine, placenta, and amniotic fluid), dams (i.e.: A and B) and foetuses were considered for microbiota profiling.

The analysis of samples biodiversity (α-diversity) showed clustering according to dam and foetus rather than analysed tissue. Faith’s phylogenetic diversity (Fig. 2A), measured based on distances, and observed species metrics showed a significant separation dependent on foetus (Mann-Whitney test: p=0.016 and 0.019, respectively). Intra- and inter-group unweighted Unifrac distances confirmed foetus-dependent microbial profiles (p=0.01, Fig. 2B). Both the metrics showed a separation dependent also on dam (permutation based test: p=0.004 and 0.036, respectively). Interestingly, no significant separation was observed based on tissue type (p>0.05), independently from the metric used to compare distributions.

**Figure 2.**
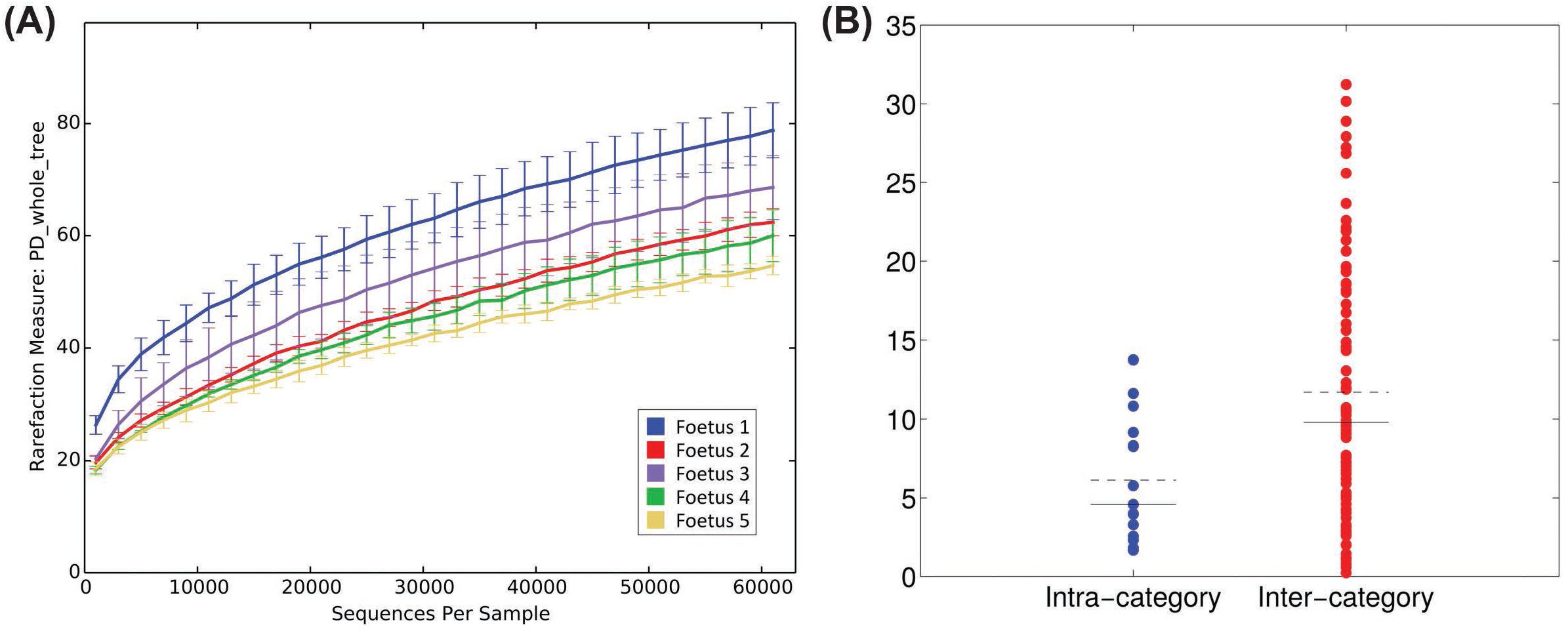
Microbial biodiversity (α-diversity) is foetus-specific. **(A)**α-diversity rarefaction curves according to Faith’s phylogenetic distance (“PD whole tree”). X-axis reports the number of sequences per sample, whereas Y-axis shows the value of the metric. Samples are grouped based on foetus number. **(B)** Distribution of distances between α - diversity PD whole tree values; distances are labelled as “intra-“ or “inter-category” according to foetus number. Dashed black line represents the mean of the distances, whereas the solid black line represents the median.

To evaluate whether different samples were characterized by distinct microbiota composition profiles (β-diversity, Fig. 3A), the distribution of Unifrac distances was assessed. As with the α-diversity, the β-diversity analyses clustered according to foetus (adonis test p=0.009 and 0.006 on unweighted and weighted Unifrac distances, respectively) and to dam (adonis test p=0.015 on weighted Unifrac distances).

**Figure 3.**
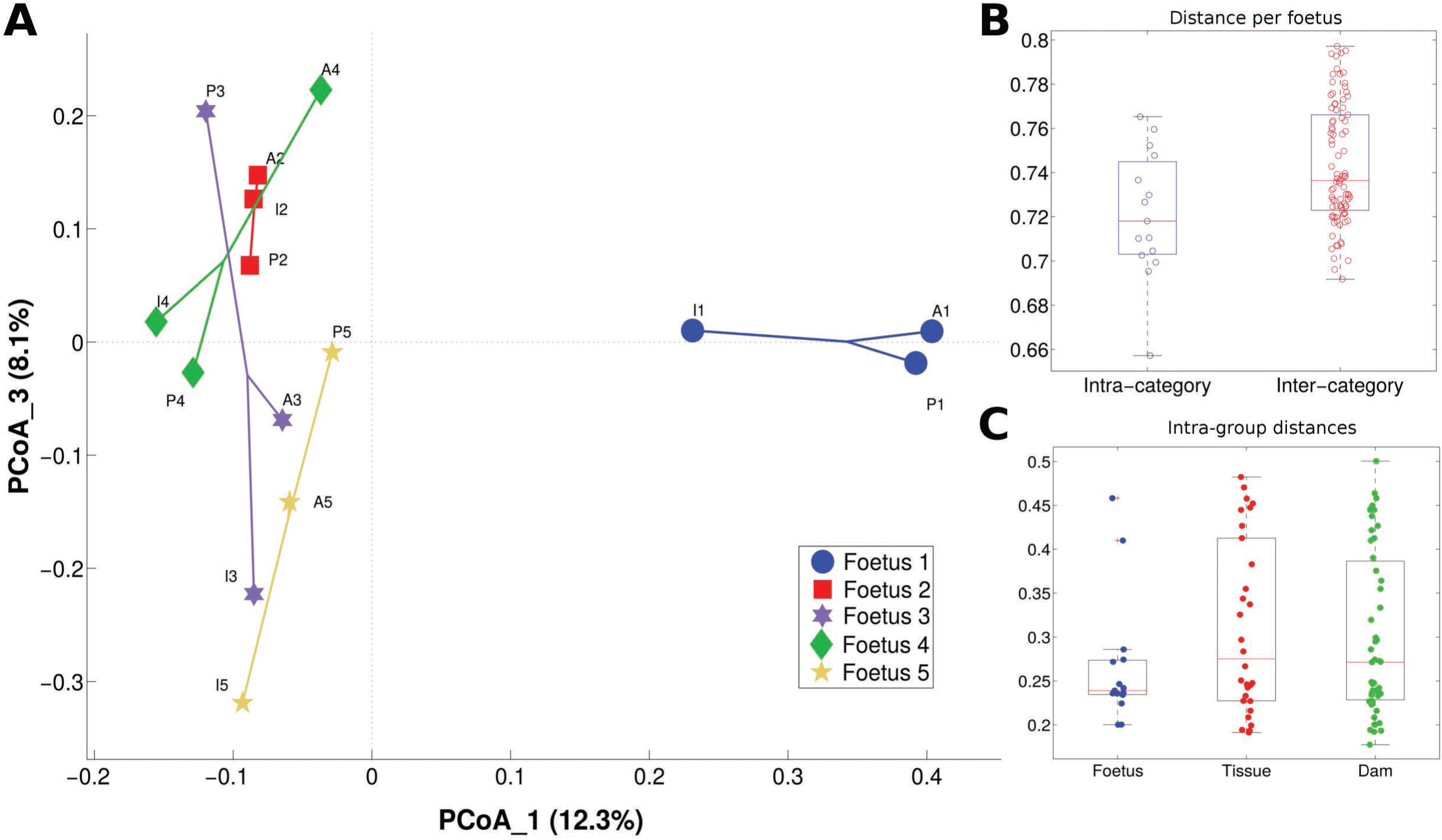
Microbiota composition (β-diversity) is foetus-specific. **(A)** Principal coordinates analysis (PCoA) of the unweighted Unifrac distances; PCoA components 1 and 3 are reported. Samples are connected together on the basis of foetus number. **(B)** Boxplots of intra- and inter-category unweighted Unifrac distances among samples; categories are based on the foetus number. **(C)** Boxplots of intra-category weighted Unifrac distances among samples; samples are grouped according to foetus, tissue or dam.

Distributions of unweighted Unifrac distances (Fig. 3B) on foetus were statistically different (p=0.01), whereas weighted Unifrac distances were not (p=0.09), indicating that significant differences are present in sub-dominant components of the microbiota. In detail, foetus 1 was characterized by a high relative abundance of *Propionibacteriaceae* (14.3% compared to an average of 1.1% in other foetuses) and *Corynebacteriaceae* (4.1% compared to an average of 0. 4% in other foetuses). Foetus 3 showed an increased abundance of *Erysipelotrichaceae* (1.7% compared to an average of 0.9% in other foetuses), and foetus 5 was enriched in *Porphyromonadaceae* (average relative abundance 2.4% compared to 1.0% in other foetuses) (Supplementary Fig. 1 and Supplementary Fig. 2A).

Analysis of distances based on sample origin (i.e. foetus, tissue and dam) showed a trend indicating similarity in microbial profiles among foetuses (Fig. 3C).

Analysis of observed species metric (p=0.036) and Faith’s phylogenetic distances (p= 0.004) allowed for a significant separation among dams, with dam B presenting with a lower biodiversity (Supplementary Fig. 3).

The microbiota signature for each tissue appeared less distinct, with only some hints of a lower presence of *Verrucomicrobiaceae* in placenta tissue (relative abundance: 1.1% compared to 2.7% and 3.7% in amniotic fluid and intestine, respectively) and a trend of higher relative abundance of *Barnesiellaceae* in the intestine (4.0% compared to 1.4% and 1.5% in placenta and amniotic liquid, respectively) (Supplementary Fig. 2C and Supplementary Fig. 4). Few differences emerged in the comparison between dam A and B (Supplementary Fig. 2B and Supplementary Fig. 5).

### Bacteria are visualised in the gut during rodent foetal development

To assess bacteria distribution, *in situ* analysis was performed on whole sectioned foetuses. Fluorescent detection revealed the presence of bacteria in the gut lumen of developing rat foetuses (Fig. 4A and 4B). In particular, eubacteria (green fluorescence) could be visualised on the different analysed sections, confirming that bacteria colonize the rodent intestine before birth. Probe for *Staphyloccaceae* did not give positive fluorescent signal in any of the tissue sections analysed.

**Figure 4.**
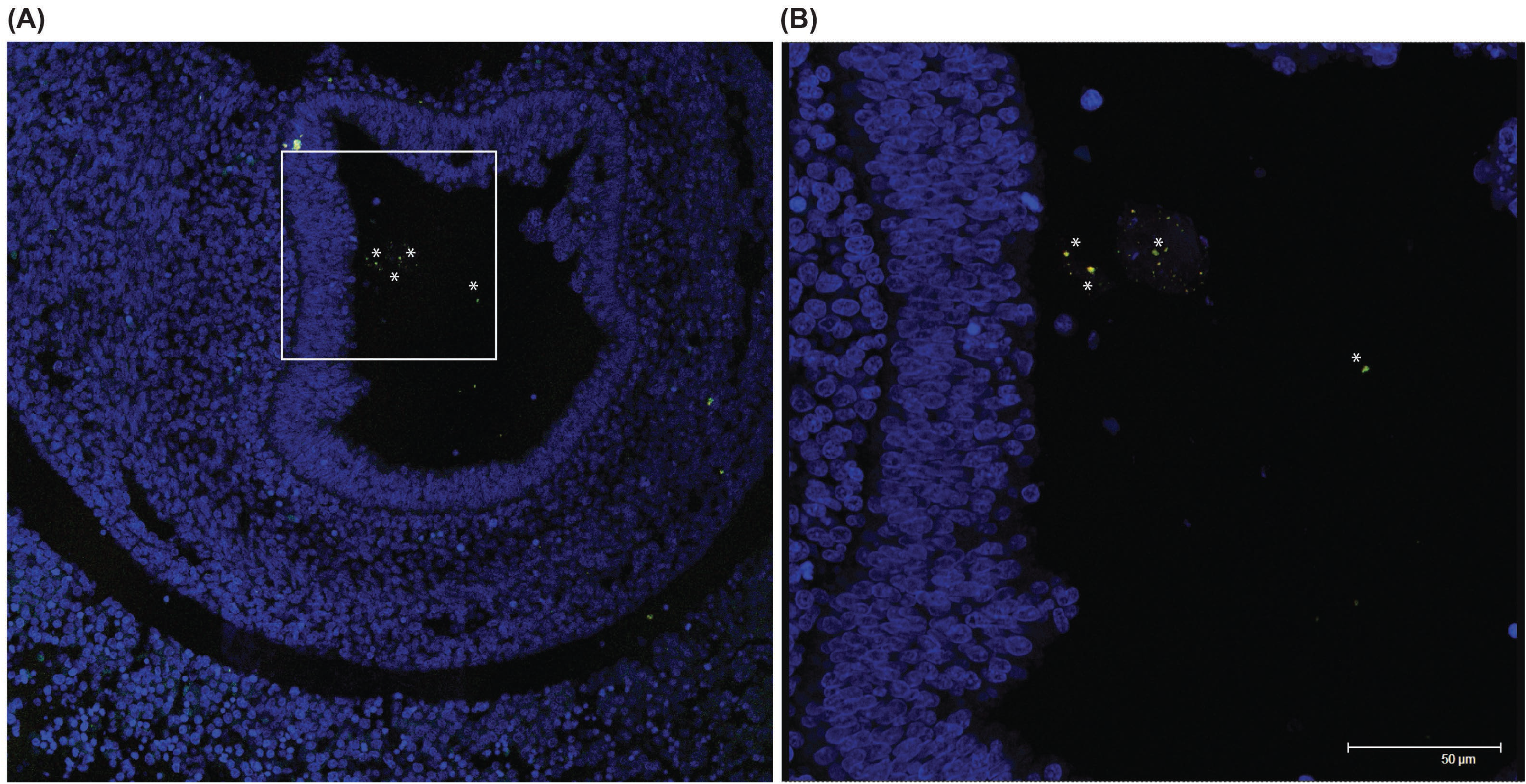
Eubacteria in the developing rodent gut lumen. Confocal microscopy images showing **(A)** eubacteria (in green) in the lumen of a 16 days post coitum rat foetus; **(B)** at a higher magnification (inset is represented as a white box in A) note the typical bacterial morphology and it is possible to identify few *Bacteroides* spp. (yellow). In blue is DAPI (nuclei) and 50 μm scale bar is reported in B. Asterisks indicate bacterial cells.

### Bacteria are present in the gut during human foetal development

Paraffin-embedded intestinal tissues from three third trimester (gestational age 29, 31 and 33 weeks) human foetuses were screened for the presence of foetal microbiota.

In all analysed samples, bacteria were observed. The most represented phyla (Fig. 5A) were *Firmicutes* (57.3±4.5), *Bacteroidetes* (17.4±1.2), *Actinobacteria* (16.8±5.4), *Proteobacteria* (4.9±2.2), and *Verrucomicrobia* (2.7±1.7). At family level, the most abundant taxa were: *Lachnospiraceae* (19.0±8.6), *Ruminococcaceae* (18.2±2.4), *Propionibacteriaceae* (9.1 ±4.1), *Bacteroidaceae* (8.9±2.2), *Streptococcaceae* (4.7±1.7), and *Veillonellaceae* (4.7±0.6).

**Figure 5.**
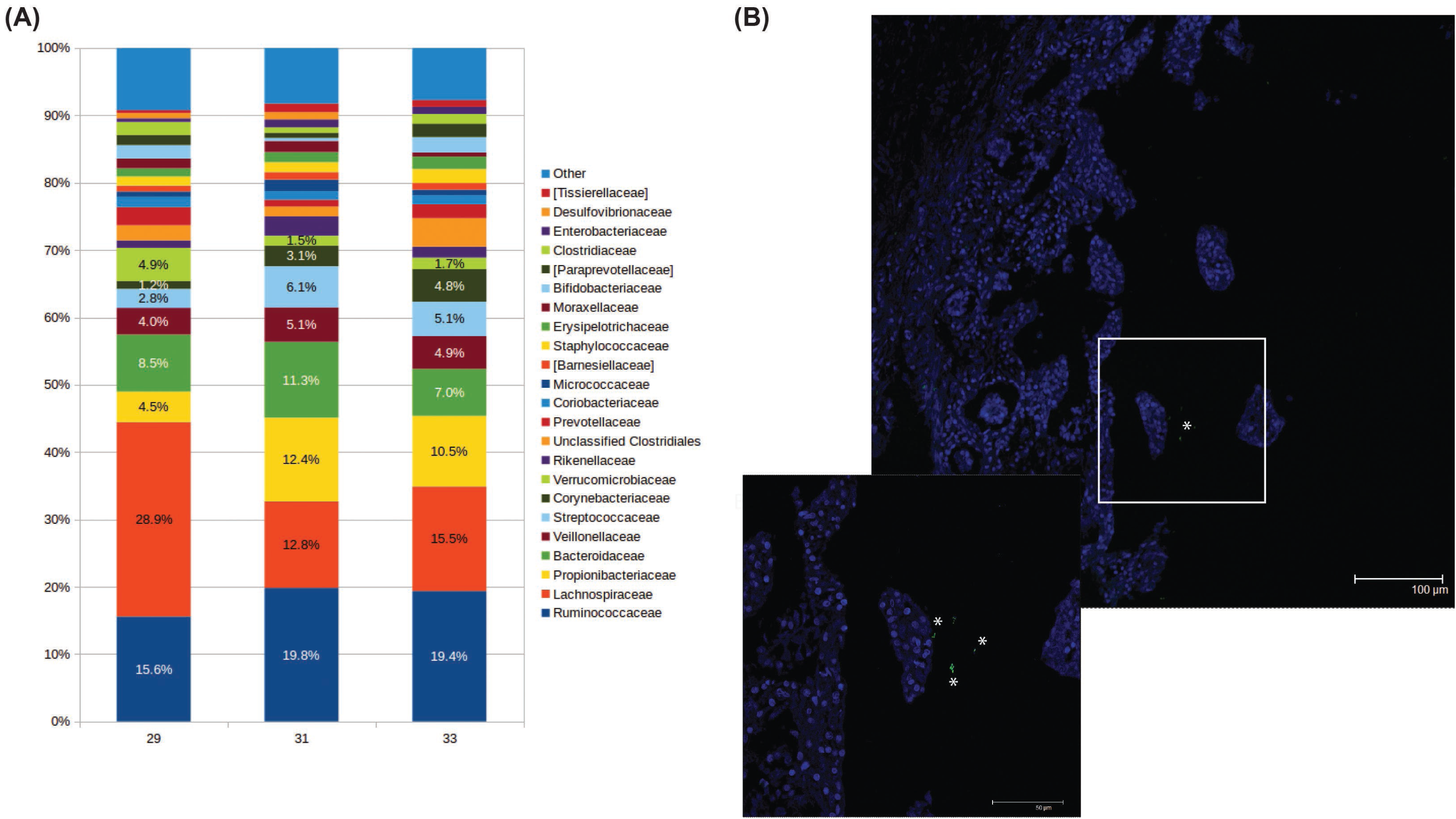
Eubacteria in the developing human gut. **(A)** Bar charts representing the family relative abundance at family level of three foetal human intestines (29, 31, and 33 weeks of gestation, respectively). **(B)** Representative confocal microscopy images of *in situ* hybridisation showing the presence of eubacteria (in green-, inset showing higher magnification) in the lumen human foetuses. In blue is DAPI (nuclei) and 100 μm and 50 μm (inset) scale bars are reported. Asterisks indicate bacterial cells.

The presence of bacteria indicated by the data obtained by NGS analysis was further validated by visualising eubacteria in the lumen of the developing gut by fluorescent *in situ* hybridization (Fig. 5B).

## DISCUSSION

Commensal microorganisms have coevolved with their natural hosts giving rise to holobionts (host plus symbiotic microbes) [13]. Microbes-encoded genes fully support and integrate the host genome in a comprehensive gene system that participates in holobiont phenotype establishment [14] and obey to fundamental genetic rules such as maternal transmission [4,15] and natural selection. Indeed, microbiota adaptation has been observed in different human niches during pregnancy [16].

Two recent papers [17,18] suggest a direct effect of maternal microbiome-immune system interaction on offspring neurodevelopment. In addition, the presence of a meconium microbiota strongly supports the existence of maternal microbial transmission *in* utero [8].

In this study, for the first time to our knowledge, we show that bacteria are present in anatomical foetal mammalian gut sections. Importantly, some species were found to be viable following specific culturing conditions. Given the limited dimension of the rat foetal intestine and the consequent paucity of the sample, culture requires subsequent steps of enrichment inevitably leading to a biased selection and a underestimation of members of the microbial community.

Reported relative abundance in meconium [6,8,19] indicate a consistent presence of *Proteobacteria,* whereas in this study, both in human and rat developing gut, *Firmicutes* were found to be more represented. These changes in microbiota may be a consequence of physiological changes that occur during birth.

Supporting the important role of the microbiota community in the developing mammalian gut, the phyla composition found in analysed samples *(Firmicutes, Bacteroidetes, Actinobacteria, Proteobacteria* and *Verrucomicrobia)* closely resembles those reported in healthy adult human gut [20,21]. It is important to note that, compared to adult and childhood tissues [22,23], the developing gut is enriched in *Actinobacteria,* and depleted in *Bacteroidetes. Actinobacteria* abundance has been shown to progressively increase in infant faeces during lactation and to then decrease when solid food is introduced in the diet. The opposite has been reported for Bacteroidetes [24-26], Hence, the difference between foetal and adult gut composition is in line with infant faecal microbiota, suggesting that solid food might be responsible for the switch. It could also suggest that the presence of a maternally provided reservoir of bifidobacteria, that with human milk oligosaccharides are known to be fundamental for the development of a balanced infant microbiota and a fully functional gastrointestinal tract [24,27-29].

Another difference found in our study compared to bacteria composition reported in meconium collected after birth [6,8], is that *Staphylococcaceae* and *Streptococcaceae* appear to be less abundant during uterine life. It is known that staphylococci are characteristic of higher respiratory tract and skin microbiota, hence it is conceivable that colonisation occurs during birth or from the first days of life through contact with maternal tissues. It has also recently been shown that both genera are abundant in colostrum and maternal milk [30], indicating a possible dual colonising path.

Analysis of 16S rRNA amplicon data clearly showed that microbiota composition is foetus- and dam-specific rather than tissue specific. Indeed, bacterial families found in amniotic fluid and placenta overlap with those found in the corresponding foetus. Importantly, the specificity seems to be independent of growing environment (i.e., uterine tissues etc.) but it seems to relate to micro niches (i.e. embryonic implant). This is true not only for the microbial community, but also for other developmental determinants, such as genetic, metabolic, biochemical or epigenetic components known to be specific to the single developing organism and not always shared among all siblings. This is in addition to the possible confounding factors of gender, time and site of implantation etc.

Given that bacteria colonise mammalian gut during intrauterine life, the fundamental question remains as to the source and the path of this in utero seeding or exposure. Recent findings [3] have described how, analysing a significant number of placentas collected after birth, the placental microbiota shares more similarities with that of the oral composition compared to vaginal, skin and/or gut communities. Rather than identifying the origin of placental microbiota, these data support the exclusion of a passive dispersion through excreting organs. In the literature, a possible microbiota colonisation *in utero* has been often hypothesised as a consequence of pathogens known to be able to reach the developing foetus. Our data suggest an alternative mechanism, where pathogens may pass the maternal barrier as a consequence of the necessary permissiveness to commensal bacteria [4], instead of resulting from infectious events (reviewed in Doran *et al.* [31])

Clearly, considering the accumulating evidence for a strict relationship between microbiome and health status, in all studied settings (age, gender, ethnicity, mtDNA SNP, haplogroup, etc.) [32-37] and the importance of maintaining or replenishing the microbial community in pathological conditions [35,38,39], the present paper opens an enormous field of investigation relating to the management of healthy pregnancies [40]. To pave this way, basic questions should first be addressed, as when colonisation occurs and through which routes, and improvement in culturomics might in the future allow for establishing *in vitro* foetal microbiota studies [41].

In conclusion, through detecting bacteria in rat and human developing gut both by Next Generation Sequencing and by *in situ* visualisation, we provide evidence that mammals do not, as often postulated, grow to be super-organisms [42-44], but instead they are born as such.

## METHODS

### Animals and housing

CD albino rats (Sprague-Dawley) were maintained in standard conditions (light 6 a.m.-6p.m., T=22+/-2°C, humidity = 55+/-5%) with tap water and food (Mucedola standard diet) *ad libitum.* Virgin females were caged overnight with males of proven fertility. The day of positive vaginal smear was considered as day 0. The pregnant rats were housed individually. Animals were euthanized by carbon dioxide inhalation and bilaterally pregnant uterine samples were collected in the morning of gestational day 16. Amniotic fluid, placentae and foetal intestines were dissected, collected in sterile conditions, and stored at -80°C until use. For dissection, uteruses were placed in sterile saline solution under laminar flow cabinet, and all procedures were conducted in the hood using UV sterilised equipment within a sterile field created by a Bunsen burner, until samples were placed in sterile tubes. For *in situ* hybridisation, whole foetuses were collected in paraformaldehyde 4% (v/v), and kept at 4°C in rotation for 4 days. Samples were then washed twice in phosphate saline solution (PBS), rehydrated through a graded series of alcohols, and paraffin-embedded.

All animal procedures were conducted in accordance with the ethical guidelines approved by the University of Milan in compliance with national (Dlgs 26/2014) and international laws and policies (EEC Council Directive 86/609).

### Human samples

From a database of foetal autopsies performed after legal termination of pregnancy, we selected, among cases without malformations and with normal karyotype, three random foetuses on the basis of gestational age, and of gut tissue availability. 4 μm thick tissue sections from formalin-fixed, paraffin-embedded tissue's samples were cut and processed for *in situ* hybridization by deparaffinisation in xylene and rehydration through a graded series of alcohols. For NGS analysis, tissue sections, cut under laminar flow cabinet using sterile blades and placed in sterile tubes, were washed twice with 1 mL of Histo-Clear (Sigma Aldrich, Milan, Italy) for 15 minutes with rotation at 56°C or until diaphanisation. Tissue was recovered by centrifugation, washed in ethanol and dried for DNA extraction.

The study was exempt from Institutional Review Board approval because, following Italian Data Protection Act 9/2013, autopsy material sampled for diagnostic purposes can be used for research as long as patient privacy is ensured. This law is in line with European Commission recommendation n. Rec(2006)4.

### Next Generation Sequencing analysis

Total bacterial DNA extraction was performed using the QIAamp DNA Microbiome Kit (QIAGEN, Hilden, Germany), following manufacturer’s instructions. Particular attention was paid to avoid environmental contamination of collected samples, and cross-contamination between samples. Samples were individually processed for DNA extraction under laminar flow cabinet, following UV sterilisation.

16S rRNA gene amplicon libraries were performed with a two-step barcoding approach according to Illumina 16S Metagenomic Sequencing Library Preparation (Illumina, San Diego, CA, USA), which amplifies two hypervariable regions (i.e.: V3, V4) of the 16S rRNA bacterial gene. Library concentration and exact product size were measured using a KAPA Library Quantification Kit (Kapa Biosystems, Woburn, MA, USA) and Agilent 2100 Bioanalyzer System (Agilent, Santa Clara, CA, USA), respectively. Agilent analysis for evaluating the correct predicted size of amplicons showed no bands in negative controls, extracted and processed in parallel with samples (Supplementary Fig. 6). Prior to sequence, libraries were pooled using Amicon Ultra 0.5ml Centrifugal Filters (Merck Millipore Ltd, Tullagreen, Carrigtwohill Co, Cork, Ireland).

The resulting library was loaded on a MiSeq^®^ 500 cycle-v2 cartridge to obtain a paired-end 2x250 bp sequencing. Demultiplexed FASTQ files were generated by Illumina MiSeq Reporter and 2.5 Gbases were obtained.

### Fluorescence *in situ* hybridization (FISH) analysis

Paraffin-embedded tissue specimens (4 μm thick) were deparaffinised by sequential steps in xylene. Then samples were re-hydrated in 95% ethanol, 90% ethanol, and finally deionized water. The slides were air-dried prior to hybridization.

FISH probe sequences for Eubacteria (EUB 338-I, 5’-GCTGCCTCCCGTAGGAGT-3’; EUB 338-III, 5’-GCTGCCACCCGTAGGTGT-3’), encompassing all bacterial species in Bacteria domain (labelled with FITC), *Bacteroides* (BAC303, 5’-TCCTCCATATCTCTGCGC-3’, Cy3), and *Lachnospiraceae* (LACHNO, 5’-TTCCCATCTTTCTTGCTGGC-3’, Cy5) were obtained from probeBase website [45]. Negative control probe (complementary to EUB 338-I probe, NON-EUB, 5’-ACTCCTACGGGAGGCAGC-3’) was also hybridised to evaluate non-specific binding. *Staphylococcaceae* probe (STAPHY, 5’-TCCTCCATATCTCTGCGC-3’, Cy3) sequence was designed as described by Gey et al. [46], and used to assess possible environmental contaminations in sampling. Probes were purchased from Integrated DNA Technologies (IDT, San Jose, CA, USA). Hybridisation was carried out using standard methods [46,47]. Briefly, sections were deparaffinised and rehydrated in serial solutions. Following section air-drying, specific oligonucleotide probes were hybridised using conditions optimized for each probe for stringent hybridisations: BAC303 at 48°C and 10% formamide; STAPHY, and LACHNO at 48°C and 30% formamide; EUB 338-I, and EUB 338-III, at 48°C and 10% or 30% formamide according to the paired probes.

DAPI (4′,6-diamidino-2-phenylindole) counterstaining was applied to assess prokaryotic and eukaryotic nuclear morphology.

In this set of experiments, a number of controls were used: sections hybridised with STAPHY probe, that resulted negative, to exclude common contaminants; sections hybridised with NON-EUB probe, as negative control; artificially contaminated sections hybridised with STAPHY probe, that resulted positive, as technical control.

Images of probe-labelled sections were acquired using a confocal laser scanning microscope (CLSM, TCS SP2, Leica, Wetzlar, Germany). Microorganisms were checked for position, size and morphology. Confocal images were acquired by series and sequential scan mode. Photomultiplier tube detectors were adjusted to minimise the bleed-through of fluorescent emissions and to optimise signal/noise ratio, in particular versus tissue auto fluorescence.

### Culturing of rodent intestine samples

A pool of dissected foetal intestines, previously frozen at −80°C, were homogenised in 1 mL of brain-heart infusion (BHI, supplemented with L-cysteine and 1 pg/ml rezasurin) broth and incubated for 3 days at 37°C in anaerobic conditions by using an anaerobic jar and an atmosphere generator (BD Diagnostics, Heidelberg, Germany). Following the pre-incubation step, Gram staining was performed.To further promote the growth of anaerobic bacteria, we inoculated the pre-incubated broth in an anaerobic blood culture bottle (BD Bactec Plus Lytic/10 Anaerobic, Germany). Finally a subculture step, using both microaerophilic and anaerobic conditions, was performed on Columbia CNA agar, TSA II + 5% Sheep Blood, MacConkey, chocolate and BHI agar. Colony identification was performed, after 48h incubation, using mass spectrometry (MALDI-TOF MS, Biomérieux).

### Data analysis

Sequencing reads were processed, filtered and analysed following similar procedures described in Borghi et al. [48]. Briefly, read pairs were merged together by PandaSeq software [49] discarding fragments of length <300 bases or >900 bases, as well as non-overlapping sequences. Then, fragments were quality-filtered, clustered into OTUs (Operational Taxonomic Units) and taxonomically classified against the 13.8 release of the Greengenes bacterial 16S rRNA database (http://greengenes.lbl.gov) using the QIIME suite (v. 1.8.0) [50].

Biodiversity (*α* -diversity) was evaluated by permutation-based t-tests, whereas “adonis” of the R package “vegan” was used for bacterial composition ( *β* -diversity). In addition, due to the reduced number of samples per category, we devised an alternative strategy for comparing the distributions of distances within and between each experimental group for both α-diversity and β-diversity evaluations. Each sample was assigned to an experimental group according to one of the associated labels (i.e.: tissue type, dam or foetus number); then, a distance between each sample and all the others was calculated. This allowed distinguishing distances between samples belonging to the same (“intra-category” distance) or to a different (“inter-category” distance) experimental group. This strategy was applied for evaluating the absolute difference for α-diversity indexes (i.e.: chao1, Shannon index, observed species and Faith’s phylogenetic distance) and the weighted or unweighted Unifrac distances [51] (β-diversity). A Mann-Whitney U-test was applied for comparing the distributions of “intra-” and “inter-category” distances.

Details of statistical methods are provided in Supplementary information.

Data availability. Raw sequence data determined in this study are available at NCBI Short Read Archive (SRA, https://www.ncbi.nlm.nih.gov/sra/) under Accession numbers PRJNA379373 and PRJNA379370.

## ACKNOWLEDGMENTS

This study was funded by a Research Grant 2016 of the European Society of Clinical Microbiology and Infectious Diseases (ESCMID) to F.B, and by Università degli Studi di Milano. We thank Raffaella Adami for her assistance in confocal laser microscopy techniques, Francesca Di Renzo, Silvia Rigamonti, and Caterina Biassoni for technical assistance. We are grateful to Ms Dawn Savery, Dr Jon Wilson and Dr Nicola Segata for commenting on the manuscript.

## AUTHOR CONTRIBUTIONS

E.B., V.M. and F.B. conceived and designed the study; E.M and V.M. performed rodent housing and sampling; L.A. and G.B. performed human tissue sampling; E. B. F.B. and G.F. performed experiments; M.S. performed bioinformatics analysis; M.S., E.B. and F.B. analysed the data; V.M. and E.B. wrote the paper. All authors made comments on the manuscript; G.M. supervised the project.

